# Offset-related brain activity in the ventrolateral prefrontal cortex promotes long-term memory formation of verbal events

**DOI:** 10.1101/828855

**Authors:** Angela Medvedeva, Rebecca Saw, Miroslav Sirota, Giorgio Fuggetta, Giulia Galli

## Abstract

Recent evidence suggests that brain activity following the offset of a stimulus during encoding contributes to long-term memory formation, however the exact mechanisms underlying offset-related encoding are still unclear. Here we used repetitive transcranial magnetic stimulation (rTMS) to investigate offset-related activity in the left ventrolateral prefrontal cortex (VLPFC). rTMS was administered at different points in time around stimulus offset while male and female participants encoded visually-presented words (first rTMS experiment) or pairs of words (second rTMS experiment) and the analyses focused on the effects of the stimulation on subsequent memory performance. The results show that rTMS administered at the offset of the stimuli, but not during online encoding, disrupted subsequent memory performance. In the first experiment we show that rTMS specifically disrupted encoding mechanisms initiated by the offset of the stimuli rather than general, post-stimulus processes. In the second experiment, we show a robust decline in associative memory performance when rTMS was delivered at the offset of the word pairs, suggesting that offset-related encoding may contribute to the binding of information into an episodic memory trace. A meta-analysis conducted on the two studies and on a previously published dataset confirmed that the involvement of the left VLPFC in memory formation is initiated by the offset of the stimulus. The offset of the stimulus may represent an event boundary that promotes the reinstatement of the previously experienced event and episodic binding.

**SIGNIFICANCE STATEMENT:** How well an event is encoded predicts how well it is remembered, and verbal encoding is an important part of everyday memory that, if disrupted, can lead to difficulties and disorders. The timing of encoding processes relative to the presentation of an event is important for successful retrieval, and little is known about the interval immediately after an event’s presentation (post-stimulus offset) which is thought to involve critical encoding processes in the VLPFC and hippocampus. The current studies demonstrate that indeed, verbal encoding processes in the VLPFC that are necessary for memory formation are triggered by the offset of the word, and these processes may involve VLPFC-hippocampal interactions that promote binding of event features into a single, coherent memory trace.

## INTRODUCTION

The type of brain activity engaged when individuals are exposed to new information is crucial to determine whether that information will be later remembered. Although research has traditionally focused on neural processes occurring online during the presentation of an encoding event, recent studies have demonstrated that peri-encoding activity –brain activity immediately preceding or following encoding– is also relevant to long-term memory formation (Cohen et al., 2015). On the one hand, EEG and fMRI research has demonstrated that brain mechanisms occurring during the anticipation of to-be-remembered information can predict subsequent retrieval (Otten et al., 2006; Park and Rugg, 2010; Galli et al., 2011, 2012, 2013, 2014). On the other, a separate line of research revealed that immediate post-stimulus processes are also critical for memory encoding (Ben-Yakov and Dudai, 2011; Rossi et al., 2011; Ben-Yakov et al., 2013; Galli et al., 2017). In fMRI studies, brain activity occurring within seconds after the termination of a visual stimulus in a set of regions including the hippocampus correlated with subsequent memory performance, possibly reflecting the binding of event features into cohesive episodic representations (Ben-Yakov and Dudai, 2011; Ben-Yakov et al., 2013).

fMRI, however, is not the optimal technique to examine the temporal dynamics of memory formation given the sluggish hemodynamic response and the correlational nature of fMRI data. Instead, repetitive Transcranial Magnetic Stimulation (rTMS) allows causal inferences on the necessity of targeted brain regions at given time intervals by temporarily interfering with neural activity in those regions at specific points in time. rTMS studies demonstrated that the engagement of the lateral prefrontal cortex at different points in time during encoding, including post-stimulus time windows, is necessary for memory formation (Machizawa et al., 2010; Rossi et al., 2011). Importantly, one recent study showed that rTMS administered over the left ventrolateral prefrontal cortex (VLPFC) within 100 ms of the offset of word stimuli disrupted the accuracy of retrieval in a subsequent memory test (Galli et al., 2017). This effect was not evident when the stimulation was delivered during the presentation of the words or at later points in time (i.e., 200-400 ms) after their offset. That study was not designed to examine offset-related brain activity *per se*, but the findings suggested that the offset of a stimulus plays a key role in the formation of new verbal memories by triggering encoding-related activity in the left VLPFC, which is implicated with the encoding of verbal information (for reviews, Blumenfeld and Ranganath, 2007; Kim, 2011; Galli, 2014). This is in line with recent research showing that event boundaries –and the offset of a visual stimulus can be unarguably considered as such–promote episodic memory formation by reinstating and binding the contents of the previously experienced episode (Sols et al., 2017; Silva et al., 2019).

This study aims to characterize the response of left VLPFC to the offset of verbal stimuli and its role in long-term memory formation using rTMS. In Experiment 1 we systematically varied word duration and time of rTMS administration to examine whether encoding-related brain activity in the VLPFC is specifically initiated by the offset of a visual stimulus or is rather activated during offset-invariant encoding processes occurring after the termination of the stimulus. A behavioral control experiment was performed to rule out effects of word duration on memory performance (Experiment 2). We also explored the idea that offset-related activity supports the binding of event features into an episodic representation (Ben-Yakov and Dudai, 2011; Ben-Yakov et al., 2013). To this end, in Experiment 3 we administered VLPFC rTMS during relational encoding and hypothesized that rTMS administered at the offset of word pairs would impact subsequent associative memory. The involvement of the left VLPFC in offset-related encoding relative to online encoding was further assessed with a meta-analysis conducted on the data from both experiments of the current study and on the data reported in Galli et al. (2017).

## MATERIALS AND METHODS

### Participants

Sixty-six native English speakers aged 18-30 years with normal or corrected-to-normal vision and good general health were recruited for the experiments. Of these, 24 (14 females; mean age ± SD: 21 ± 3 years; range: 18-29 years) took part in Experiment 1; 18 (15 females; mean age ± SD: 22 ± 3 years; range: 19-29 years) took part in Experiment 2; and 24 (17 females; mean age ± SD: 20 ± 1 year; range: 18-23 years) took part in Experiment 3. For the rTMS experiments (Experiments 1 and 3), we calculated that a minimum of 22 participants was required to detect an effect size of *d* = 0.74 (as in Galli et al. 2017, assuming α = 0.05 and 1-ß = 0.95, one-tailed paired-samples t-test), and adjusted the final sample size upwards to account for possible attrition rate. Two subjects in Experiment 1 and one subject in Experiment 3 were tested but excluded from the analyses due to low memory performance based on a-priori determined exclusion criteria (± 2 SD from the average). Participants received course credits or monetary compensation for their participation. All participants gave written informed consent. The studies were approved by the University of Roehampton Ethics Committee (Experiment 1 and 3) or the Kingston University Ethics Committee (Experiment 2).

### Materials

In Experiment 1 (rTMS) and Experiment 2 (behavioral control experiment), stimuli were 288 seven-letter words (mean word frequency = 23.03, SD = 39.86; Kučera and Francis, 1967) extracted from the MRC psycholinguistic database (Coltheart, 1981). For each subject, 180 words were randomly selected from this pool to be presented as old items during the study phase and 108 words served as new items in the test phase. In Experiment 3 (rTMS), a list of 720 words of three or fewer syllables was extracted from the MRC psycholinguistic database and used to create 360 word pairs matched for frequency and imageability. For each subject, 240 word pairs were randomly selected to be presented as old pairs during the study phase and the remaining 120 word pairs were used as novel word pairs. For all experiments, an additional list of words or word pairs was used as practice for the study and test tasks.

### Behavioral Task

All experiments consisted of an intentional encoding task followed by a memory task after a delay of approximately 5 minutes. Experiment 1 (rTMS) and 2 (behavioral control experiment) employed an item memory task and Experiment 3 (rTMS) employed an associative memory task.

In Experiment 1 and 2, at study participants viewed a total of 180 words, presented one at the time, and were asked to indicate whether the word was pleasant or unpleasant by pressing one of two keys on the keyboard with their right or left index finger. This task ensured that participants attended to each word and encouraged deep encoding of the stimuli. Participants were also instructed to memorize the words in view of a subsequent memory test. Trials started with a fixation mark that stayed on the screen for 1000 ms followed by the presentation of the word. In both experiments, the duration of the word varied as a function of the experimental condition (see *rTMS protocol and experimental conditions* below and Figure 1A), and the inter-trial interval was varied accordingly to achieve a trial duration of 5600 ms in all conditions. The total number of trials and the ratio of old/new words was identical in the two experiments, but the number of trials in each presentation block was different to accommodate a different number of experimental conditions. In Experiment 1, words were presented in six study blocks of 30 words each, corresponding to the six stimulation conditions. In Experiment 2, words were presented in four study blocks of 45 words each, corresponding to four word-duration conditions.

**Figure 1:**
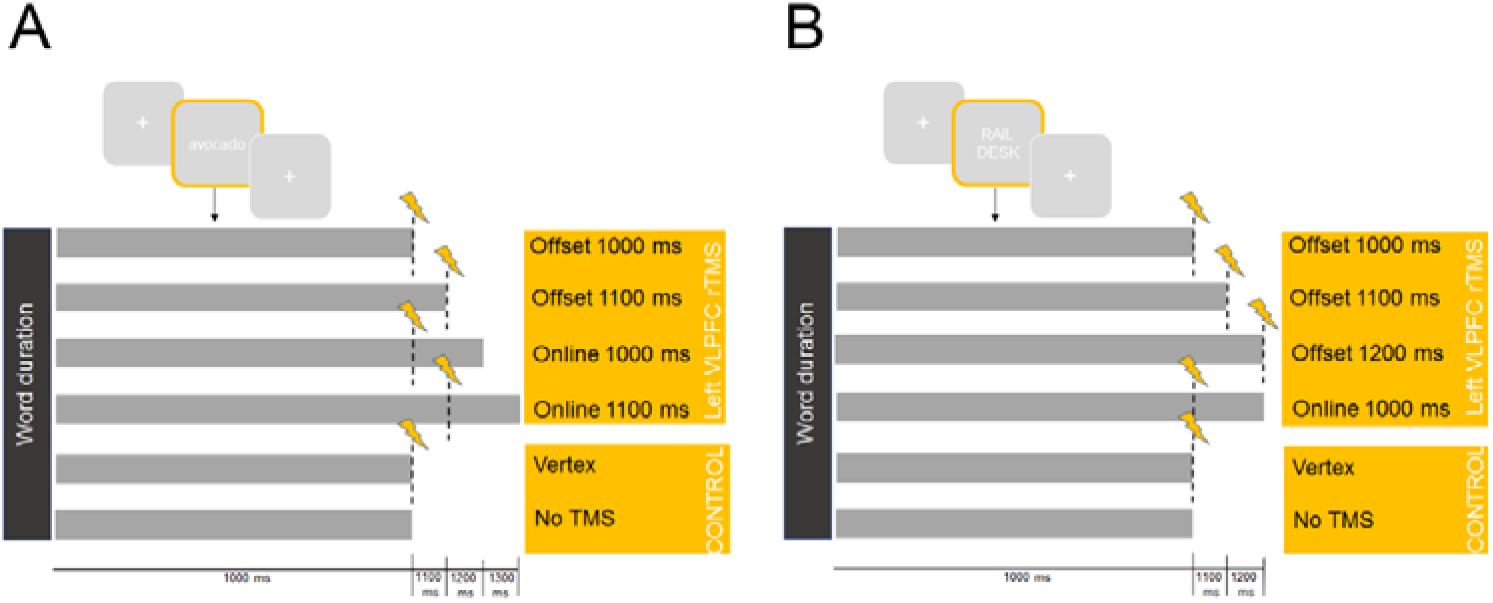
Schematic illustration of the rTMS conditions in Experiment 1 **(A)** and Experiment 3 **(B)**.

In the test phase, the words from each study block were interspersed with 18 (Experiment 1) or 27 (Experiment 2) new words and presented again for the recognition memory task, resulting in six test blocks of 48 items each (Experiment 1) or four test blocks of 72 words each (Experiment 2). The presentation of blocks in the test phase followed the same order of the study phase (e.g., words that were presented in the first block in the study phase, were presented in the first block of the test phase). For each word, participants had to decide whether or not they had seen the word during the study phase by pressing one of two keys with their right or left index fingers. The assignment of old responses to the left or right hand was counterbalanced across subjects. Each trial started with a 1000 ms fixation, followed by a word that stayed on the screen for 1000 ms. The time in between the offset of the word and the onset of the following trial was 1500 ms.

In Experiment 3, at study participants viewed a total of 240 word pairs, presented one at the time in six consecutive blocks corresponding to six stimulation conditions (see *rTMS protocol and experimental conditions* below and Figure 1B). The word pair was presented at the center of the computer screen, one word above the other. Participants were asked to create a mental image incorporating the concepts represented by both words and press one of two keys on the keyboard with their right or left index finger to indicate whether the quality of the mental image was good or bad. Participants were also instructed to memorize the word pairs and it was emphasized that the relationship between the two words of each pair was important for the following memory test. Each study trial started with the presentation of a fixation mark for 1000 ms, followed by the presentation of the study pair. As in Experiment 1 and Experiment 2, time the word pair remained on the screen varied as a function of the experimental condition, and the inter-trial interval was varied accordingly to achieve a trial duration of 5600 ms in all conditions.

In the test phase, the 40 pairs of studied items from each study block were presented again and intermixed with 20 novel word pairs, resulting in six test blocks of 60 word pairs each. Of the 40 pairs of studied items in each block, 20 remained in the same pairing as at study (‘intact’ pairs) and 20 were rearranged such that the studied items were the same but reassembled in different pairs (‘rearranged’ pairs). Presentation location (above or below) of the pairs of studied items and order of blocks was maintained between study and test. Participants were asked to decide whether the items had been paired together at study (intact judgment), presented at study but on separate trials (rearranged judgment), or not presented at study (new judgment). To ensure that associative judgements were not contaminated by guessing, participants were also instructed to respond ‘rearranged’ when uncertain if a test pair was intact (Addante et al., 2015). Intact and rearranged judgements were given by pressing one of two keys of the keyboard with the index or middle finger of one hand, whereas novel judgements were given by pressing a third key with the index finger of the other hand. The hand with which responses were made was counterbalanced across participants. Each trial started with a 1000 ms fixation, followed by a word that stayed on the screen for 1000 ms. The time in between the offset of the word and the onset of the following trial was 1500 ms. In all experiments, fixation marks and words were presented in a white uppercase Helvetica on a gray background using the Cogent 2000 toolbox (http://www.vislab.ucl.ac.uk/cogent.php).

### rTMS protocol and experimental conditions

In the rTMS experiments (Experiments 1 and 3), trains of 600 ms 20 Hz rTMS were delivered in the study phase through a MagStim Super Rapid stimulator with a biphasic current waveform (Magstim, UK). A figure-of-eight 70-mm coil was used for the stimulation. The coil was placed tangentially to the scalp, with the handle pointing backwards and laterally at a 45° angle of the middle sagittal axis of the participants’ head. Prior to rTMS, single magnetic pulses were delivered to the hand area of the left motor cortex to establish the individual excitability threshold for the first dorsal interosseous muscle (Rossini et al. 2015). For each subject, the intensity of the stimulation during the experiment was set to 90% of the individual motor threshold. The VLPFC stimulation site was identified on the scalp using a TMS-magnetic resonance imaging coregistration system (SofTaxic, Italy). The VLPFC coordinates were automatically estimated by the Navigator System, on the basis of an MRI-constructed stereotaxic template. MNI coordinates for the left VLPFC (−53, 28, 12) corresponded to those used in previous rTMS studies of subsequent memory effects (Blumenfeld et al., 2014).

Figure 1 illustrates the experimental conditions in the two rTMS experiments. All experiments followed a repeated measures design with rTMS condition as the within-subjects factor. In our previous work (Galli et al., 2017) the time of rTMS administration varied while the duration of word stimuli was set to 1000 ms in all conditions. As a consequence, it was not possible to ascertain whether rTMS effects observed immediately after word offset were driven by the offset itself, or by offset-invariant post-stimulus processes occurring around 1000 ms after the onset of the words, therefore temporally coinciding with word offset. To adjudicate between these two competing explanations, in Experiment 1 testing item memory we systematically varied word duration and time of rTMS administration. We used four VLPFC stimulation conditions, corresponding to four combinations of word duration/time of rTMS onset (Figure 1A). In the *Offset 1000 ms* and *Offset 1100 ms* conditions, rTMS was delivered at the offset of 1000- and 1100-ms words respectively. In the two online conditions, rTMS was delivered at the same timing of the offset conditions but while the words were still on the screen. More specifically, in the *Online 1000 ms* condition rTMS was delivered 1000 ms after the onset of 1200-ms words. In the *Online 1100 ms* condition rTMS was delivered 1100 ms after the onset of 1300-ms words.

In Experiment 3 testing associative memory, three VLPFC stimulation conditions were identical to Experiment 1 (*Offset 1000 ms, Offset 1100 ms* and *Online 1000 ms*), with the exception that we removed the *Online 1100 ms* and added an additional offset condition (*Offset 1200 ms*) to investigate offset-related effects at later temporal windows.

In both experiments, participants additionally received vertex stimulation and performed one block of the task without rTMS, which made six experimental conditions in total (Figure 1). The vertex stimulation site was defined as a point midway between the inion and the nasion and equidistant from the left and right intertragal notches. Since this region is not involved in learning and memory processes, it was considered a control site for possible unspecific somatosensory, acoustic, or arousal effects of active TMS (Rossi et al., 2011). In both experiments, the four PFC conditions were administered in succession to avoid coil dispositioning and their order was randomized for each participant. The order of the PFC, Vertex and No-rTMS conditions was counterbalanced in a balanced Latin Square design.

It has been previously reported that stimulation of the lateral aspects of the prefrontal cortex may be uncomfortable to some participants (Machizawa et al., 2010). Before the start of the experiments, we delivered trains of rTMS to the targeted locations and encouraged participants to report any excessive distress in order to ensure that all participants were comfortable with the TMS stimulation. Eight participants in Experiment 1 and 12 in Experiment 3 reported excessive discomfort and did not continue with the experiment. To assess the effect of any discomfort in the participants who completed the experiments, we administered a TMS sensation screening questionnaire (adapted from Rossi et al., 2011). No stimulation was delivered in Experiment 2.

### Statistical Analyses

Recognition accuracy in Experiments 1 and 2 was established with the discrimination index *Pr* (the proportion of hits minus the proportions of false alarms; Snodgrass and Corwin, 1988). Analysis of memory accuracy in Experiment 3 focused on associative hits as index of associative memory performance (‘intact’ responses to intact word pairs). Furthermore, in the three experiments we analyzed response times (RTs) at study and test. In Experiment 1 and Experiment 2 RTs for correct memory judgments were analyzed by averaging RTs for hits and correct rejections.

In both rTMS experiments, the effects of rTMS at encoding on subsequent memory accuracy and RTs was investigated by comparing each VLPFC condition with the control conditions, using Bonferroni-corrected, one-tailed (for RTs and memory accuracy in the test phase) or two-tailed (for encoding RTs) pairwise comparisons. We report corrected p values throughout the manuscript. We conducted preliminary two-tailed pairwise comparisons to examine differences between the *Vertex* and the *No TMS* control conditions and observed no significant difference in all analyses (*p*s > 0.065, see Table 1 for memory performance separately for the two conditions). We therefore collapsed the two conditions to reduce the number of comparisons and achieve a unitary baseline (Galli et al., 2017).

**Table 1:** Memory performance in Experiment 1 and Experiment 3. DI: Discrimination Index Pr (proportion of hits minus proportion of false alarms, Snodgrass and Corwin, 1988). Associative Hits: ‘intact’ responses to intact pairs. Standard deviations are displayed in parentheses.

| Experiment 1 |  |  |  | Experiment 3 |
| --- | --- | --- | --- | --- |
|  | Hits | False Alarms | DI | Associative Hits |
| Offset 1000 ms | 0.81 (0.12) | 0.24 (0.15) | 0.57 (0.18) | 0.48 (0.16) |
| Offset 11000 ms | 0.77 (0.13) | 0.26 (0.14) | 0.51 (0.22) | 0.50 (0.19) |
| Offset 1200 ms |  |  |  | 0.42 (0.16) |
| Online 1000 ms | 0.81 (0.12) | 0.27 (0.18) | 0.54 (0.19) | 0.51 (0.21) |
| Online 1100 ms | 0.79 (0.17) | 0.23 (0.14) | 0.56 (0.22) |  |
| Vertex | 0.83 (0.13) | 0.20 (0.16) | 0.63 (0.20) | 0.48 (0.15) |
| No TMS | 0.82 (0.14) | 0.26 (0.16) | 0.57 (0.23) | 0.54 (0.16) |

Experiment 2 was conducted to rule out any effect of word duration *per se* during encoding on subsequent memory retrieval. One might speculate that memory performance decreases with shorter exposure times at encoding and that any detrimental effects of rTMS arisen with shorter word durations were due to the fact that the short word duration made this condition inherently more difficult and/or more susceptible to the effects of rTMS. Therefore in Experiment 2 we compared the memory performance for the different word durations at encoding from Experiment 1 (1000 ms, 1100 ms, 1200 ms and 1300 ms) using a one-way repeated-measures ANOVA.

Finally, we meta-analyzed the data from the two rTMS experiments reported here and from Galli et al. (2017) to further test the robustness of offset-related rTMS effects and test the validity of the conclusion that, across the three experiments, rTMS induced larger effects when delivered at the offset of the words compared to online encoding. We compared the means aggregated across the collapsed online conditions (vs. control) and offset conditions (vs. control) for the discrimination index *Pr* and hit rates. We first calculated standardized mean change measures for individual studies between online/offset and control conditions and then meta-analyzed them using a random model (restricted maximum likelihood estimator) as implemented in ‘metafor’ R package (Viechtbauer, 2010). We then used time (online/offset conditions) as a moderator variable.

Statistical analyses were performed in SPSS (version 24, IBM) and R version 3.5.3 (R Core Team, 2019). All data are accessible on the Open Science Framework website (https://osf.io/xev6r/).

## RESULTS

### Memory accuracy

In Experiment 1 we found that the administration of VLPFC rTMS at the offset of 1100-ms words impaired subsequent item memory performance (*t*_*21*_ = 2.79, *p* = 0.020, *d* = 0.49; Figure 2). We did not observe any impairment of memory performance in the *Offset 1000 ms* condition and in the two online conditions (*p*s > 0.064; Table 1). Experiment 2 revealed no effect of word duration during encoding on memory accuracy (*p* = 0.814), confirming that the detrimental effects of rTMS in Experiment 1 were not due to an effect of word duration. In Experiment 3 rTMS disrupted subsequent associative memory performance when administered at the offset of 1200-ms word pairs (*t*_*22*_ = 2.99, *p* = 0.014, *d* = 0.57; Figure 3). There was no decrease in associative memory in the other conditions including the online condition (*p*s > 0.164; Table 1).

**Figure 2:**
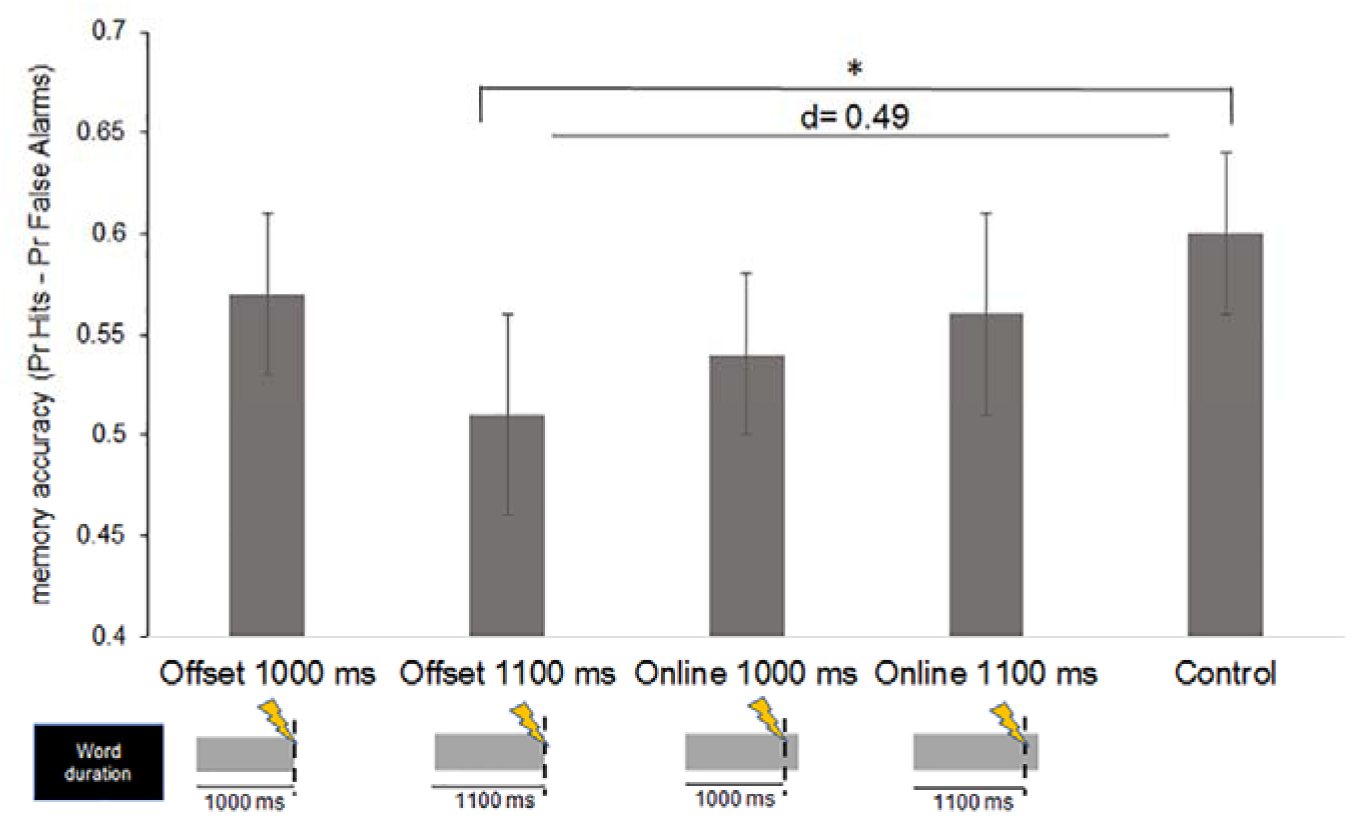
Memory performance as a function of rTMS administration in Experiment 1. A decrease in memory accuracy is evident when the stimulation was administered at the offset of 1100-ms words. The baseline (far right column) is based on the collapsed vertex and no-TMS conditions. * denotes *p* < 0.05 (Bonferroni-corrected). Effect sizes are shown as Cohen’s *d*. Error bars depict standard errors.

**Figure 3:**
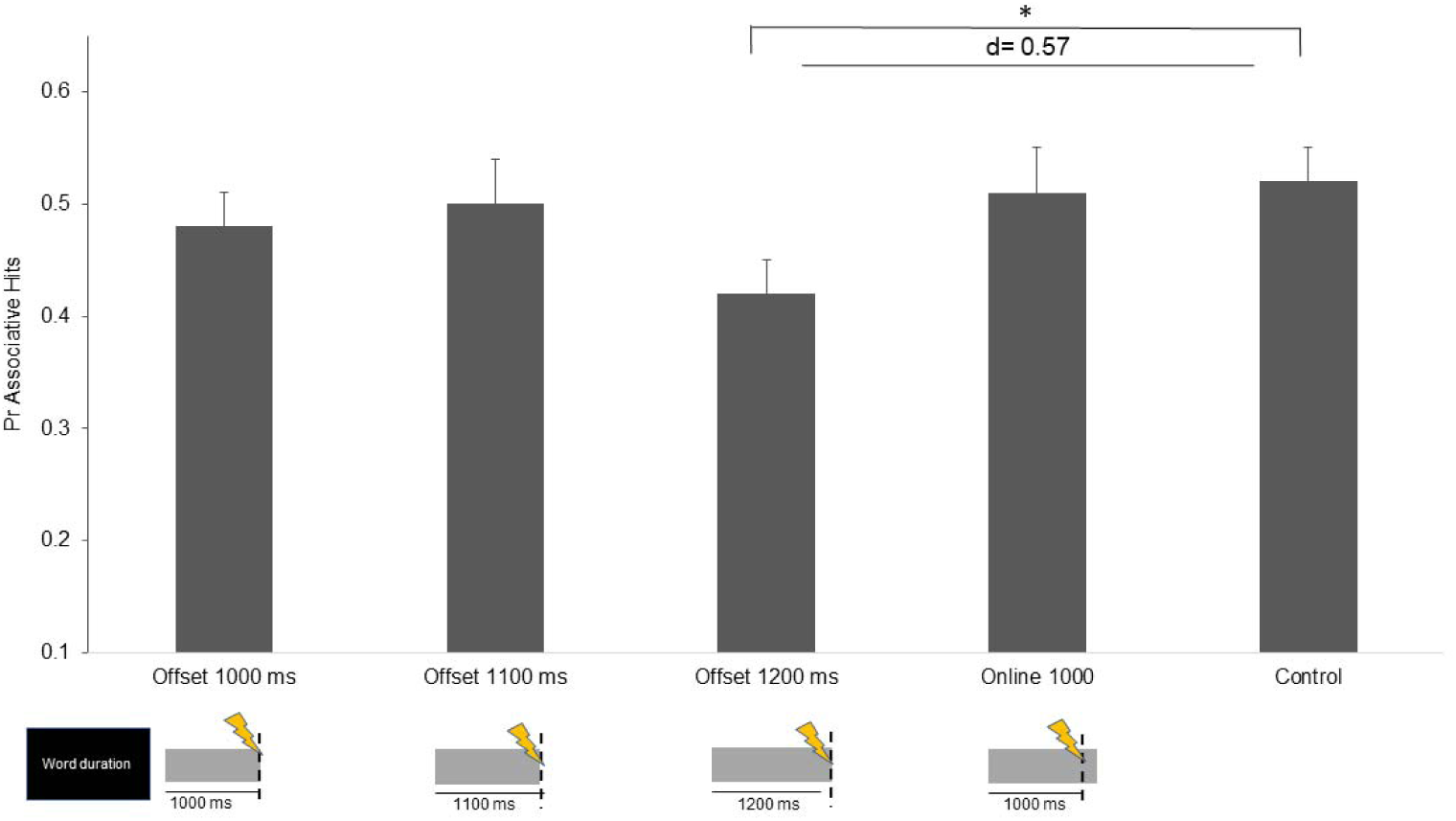
Memory performance as a function of rTMS administration in Experiment 3. Subsequent associative memory performance decreased when rTMS was delivered at the offset of 1200-ms word pairs. The baseline (far right column) is based on the collapsed vertex and no-TMS conditions. * denotes *p* < 0.05 (Bonferroni-corrected). Effect sizes are shown as Cohen’s *d*. Error bars depict standard errors.

### Reaction times

#### Encoding task

rTMS did not affect the time taken to give a response in the encoding task in either Experiment 1 (*p*s > 0.804) or Experiment 3 (*p*s > 0.084).

#### Memory task

There was no significant effect of VLPFC rTMS on test phase RTs in Experiment 1 (*p*s > 0.115), and no effect of word duration at encoding on test phase RTs in Experiment 2 (*p* = 0.280). In Experiment 3 two additional subjects were excluded from the analyses due to technical difficulties. RTs for associative hits were slower in the *Offset 1200 ms* condition (*t*_*19*_ = 2.91, *p* = 0.040, *d* = 0.66). None of the other pairwise comparisons was statistically significant (*p*s > 0.108).

### Meta-analysis

For the discrimination index, we found a small non-significant effect in the online conditions (Hedge’s *g* = 0.19, 95% CI [-0.08, 0.45], *z* = 1.38, *p* = 0.169) and a medium statistically significant effect in the offset conditions (*g* = 0.57, 95% CI [0.29, 0.86], *z* = 3.99, *p* < 0.001). We ran a moderation analysis to formally test the statistical difference between these two effects and found a significant medium-sized difference (*b* = 0.39, 95% CI [0.003, 0.78], QM(1) = 3.90, *p* = 0.048) without substantial residual heterogeneity (QE(4) = 2.99, *p* = 0.560, I^2^ = 0.0%, τ^2^ = 0). The pattern was similar for hit rates, with a small and statistically non-significant effect in the online conditions (*g* = 0.07, 95% CI [-0.19, 0.34], *z* = 0.56, *p* = 0.577) and a medium and statically significant effect in the offset conditions (*g* = 0.45, 95% CI [0.18, 0.72], *z* = 3.23, *p* = 0.001). Again, the moderation analysis showed a medium effect size difference between the two effects, but this time it did not reach statistically significance (*b* = 0.38, 95% CI [-0.002, 0.75], QM(1) = 3.79, *p* = 0.052) and without substantial residual heterogeneity (QE(4) = 1.06, *p* = 0.901, I^2^ = 0.0%, τ^2^ = 0). Note that the recommended minimal number of comparisons for a moderation analysis is typically at least ten (e.g., Borenstein, Hedges, Higgins, & Rothstein, 2009), therefore the results of the moderation analysis should be interpreted with caution.

### TMS-induced sensations questionnaire

Overall the stimulation induced discomfort in the majority of participants. Only 22% of participants in Experiment 1 and one participant in Experiment 3 reported no discomfort associated with rTMS, with the remaining participants reporting mild (27% Experiment 1, 30.5% Experiment 3), moderate (40.5% Experiment 1, 45% Experiment 3) or strong discomfort (one participant Experiment 1, 24% Experiment 3). Only a minority of participants reported that their ability to perform the task was unaffected by the discomfort (32% Experiment 1), while the remaining participants reported that the task was slightly affected (32% Experiment 1, 48% Experiment 3), much affected (23% Experiment 1, 16% Experiment 3) or considerably affected (23% Experiment 1, 36% Experiment 3). To assess whether the discomfort had any impact on the memory impairment induced by rTMS, we correlated both the intensity of the discomfort and its perceived effect on the task (in both cases coded on a scale from 1 to 4) with the significant rTMS-induced changes in Experiment 1 (*Offset 1100 ms* condition) and Experiment 3 (*Offset 1200 ms* condition). We found no significant correlation between memory impairment and general discomfort or the degree to which the discomfort affected the task (*p*s > 0.158).

## DISCUSSION

The left VLPFC has been frequently implicated with the encoding of verbal information (for reviews, Blumenfeld and Ranganath, 2007; Kim, 2011; Galli, 2014). Previous rTMS studies had demonstrated that brain activity in the left VLPFC occurring after the termination of a stimulus is critical for memory formation (Rossi et al., 2011; Galli et al., 2017). Here, in two experiments we demonstrated that the involvement of the left VLPFC during the formation of new verbal memories is specifically triggered by the offset of word stimuli. rTMS only disrupted subsequent memory performance when delivered at the offset of the words, but not during online encoding (i.e. during their presentation). This finding was confirmed by a meta-analysis that combined the data from three experiments that allowed comparison between online and offset-related activity (Experiment 1, Experiment 3 and data reported in Galli et al., 2017).

These results are broadly in line with a previous fMRI study which showed that brain activity time-locked to the offset of the stimulus in the hippocampus and the caudate nucleus was correlated with subsequent memory performance (Ben-Yakov and Dudai, 2011). That study did not reveal any engagement of the VLPFC in offset-related memory encoding, perhaps due to differences in stimulus materials (movies vs word stimuli). However, the results converge with our findings in indicating a prominent role of offset-related brain activity in memory formation. Our study using the causal approach offered by rTMS further indicates that this brain activity is not only relevant, but also necessary to memory formation. It is worth noticing that we could not examine the role of the hippocampus because the depth of TMS prevents a direct stimulation of medial temporal lobe structures. However, studies have shown that PFC stimulation can modulate network dynamics and propagate to distant brain regions, including the hippocampus (Li et al., 2004; Bilek et al., 2013). One hypothesis thus is that in the current study VLPFC stimulation at word offset interfered with memory formation through an indirect effect on medial temporal lobe structures. This idea however is at odds with the results of a recent study using intracranial EEG recordings showing that subsequent memory effects in the medial temporal lobe *precede* those found in the VLPFC (Burke et al., 2014). Further studies are needed to clarify the temporal sequence of activations during encoding across different brain regions.

What could be the specific mechanisms underlying offset-related encoding activity in the left VLPFC? The results of Experiment 3 help provide an answer to this question by showing that offset-related mechanisms may be related to the binding of information into an episodic representation. In a task that required participants to memorize the association between word pairs, we observed a large decrease in associative memory accuracy when rTMS was delivered at the offset of the pairs during encoding, along with an increase in the time taken to give memory judgments. If rTMS did not disrupt the binding of the word pairs into an integrated memory trace, we would not have observed this decrease. We speculate that the offset of a word stimulus triggers episodic binding and associative encoding processes in the left VLPFC, which in turn contribute to the formation of the memory trace. Although associative encoding is more prominent when there is a specific instruction to associate different features or items into a unique memory trace, such as in Experiment 3, it could also occur in single-item encoding in the absence of explicit associative task demands, for instance, by associating an item with the preceding ones, with previously-stored semantic information or contextual information. Therefore, episodic binding is not exclusive to associative encoding tasks, but also occurs with single item encoding as in Experiment 1 and in our previous study (Galli et al., 2017). It is notable that although associative encoding is typically associated with activity of the hippocampus (Davachi and Wagner, 2002), other studies have revealed an equally relevant role of the left VLPFC in associative memory formation (Staresina and Davachi, 2006). In addition, the left VLPFC could interact with the hippocampus due to their functional and anatomical connections (Barredo et al., 2013).

Another explanation for the current findings, which is not necessarily mutually exclusive with the interpretations above, takes into account the role of event boundaries in memory formation. Studies on sequential learning have demonstrated that memory encoding is enhanced for information presented at event boundaries, for instance when shifts in stimulus category, perceptual context or object location occur, and that these memory enhancements are related to neural activity in the hippocampus and left VLPFC (e.g., DuBrow and Davachi, 2016; Horner et al., 2017; Heusser et al, 2018). Furthermore, two recent EEG studies (Sols et al., 2017; Silva et al., 2019) found that event boundaries elicit the reinstatement of neural activity patterns associated with the just-experienced events promoting long-term memory formation. It is reasonable to assume that event offsets are experienced as event boundaries and that VLPFC activation initiated by the offset contributes to the recovery of the just-experienced events (Clewett et al., 2019). This idea is in line with the finding of similar activation patterns at the onset and immediately after the offset of visual stimuli that are successfully maintained in working memory (van de Nieuwenhuijzen et al., 2016).

One observation is that the specific time of occurrence of offset-related effects differed across experiments. rTMS effects on item memory were evident when rTMS was delivered at the offset of 1100-ms word stimuli in Experiment 1 and at the offset of 1000-ms stimuli in our previous investigation (Galli et al., 2017). Furthermore, effects on associative memory were observed when the stimulation occurred at the offset of 1200-ms word stimuli. One way to reconcile these discrepancies is to consider individual differences in encoding times. In some participants or trials, early encoding processes related to item-specific encoding (Mangels et al., 2001) may have been slower to complete, or processing times in general may have been slower. This could have resulted in a lack of rTMS effects for earlier offsets (e.g. 1000 ms in Experiment 1), especially for associative encoding, since the offset was too early to induce VLPFC activation and/or earlier encoding processes were yet to complete before associative encoding processes could initiate. Future studies could explore the relationship between offset-related brain activity and individual differences in encoding using finer-grained measures of encoding time such as EEG. Regardless of the specific differences across experiments though, the results of the meta-analysis unequivocally confirmed that memory formation was impaired when rTMS was delivered at the offset of the words, and not during their online encoding.

Finally, it is worth noting that the left VLPFC was implicated in offset-related encoding in this experiment due to its involvement in verbal memory formation (Blumenfeld and Ranganath, 2007; Kim, 2011; Galli, 2014), but more work is needed to clarify whether other brain regions are implicated using different stimulus materials.

Taken together, our findings offer insights into the temporal dynamics of memory formation and show that brain mechanisms in the left VLPFC induced by the offset of a verbal stimulus are responsible for the formation of verbal memories. By clarifying when and how memories are formed, our findings may help to refine neurorehabilitation programs in patients with memory disorders.

## Acknowledgements

This work was supported by a British Academy grant (SG170456)

## REFERENCES

Addante RJ, de Chastelaine M, Rugg, MD (2015) Pre-stimulus neural activity predicts successful encoding of inter-item associations. Neuroimage 105:21–31.

Barredo J, Öztekin I, Badre D (2013) Ventral fronto-temporal pathway supporting cognitive control of episodic memory retrieval. Cereb Cortex 25:1004–1019.

Ben-Yakov A, Dudai Y (2011) Constructing realistic engrams: Poststimulus activity of hippocampus and dorsal striatum predicts subsequent episodic memory. J Neurosci 31:9032–9042.

Ben-Yakov A, Eshel N, Dudai Y (2013) Hippocampal immediate poststimulus activity in the encoding of consecutive naturalistic episodes. J Exp Psychol Gen 142:1255–1263.

Bilek E, Schäfer A, Ochs E, Esslinger C, Zangl M, Plichta MM, Braun U, Kirsch P, Schulze TG, Rietschel M, Meyer-Lindenberg A, Tost H (2013) Application of high-frequency repetitive transcranial magnetic stimulation to the DLPFC alters human prefrontal-hippocampal functional interaction. J Neurosci 33:7050–7056.

Blumenfeld RS, Lee TG, D’Esposito M (2014). The effects of lateral prefrontal transcranial magnetic stimulation on item memory encoding. Neuropsychologia 53:197-202.

Blumenfeld RS, & Ranganath C (2007) Prefrontal cortex and long-term memory encoding: an integrative review of findings from neuropsychology and neuroimaging. Neuroscientist 13:280–291.

Borenstein, M., Hedges, L. V., Higgins, J., & Rothstein, H. R. (2009). Introduction to Meta-Analysis. Chichester, UK: Wiley.

Burke JF, Long NM, Zaghloul KA, Sharan AD, Sperling MR, Kahana MJ (2014) Human intracranial high-frequency activity maps episodic memory formation in space and time. Neuroimage 85:834–843.

Clewett D, DuBrow S, Davachi L (2019). Transcending time in the brain: How event memories are constructed from experience. Hippocampus 29:162–183.

Cohen N, Pell L, Edelson MG, Ben-Yakov A, Pine A, Dudai Y (2015) Peri-encoding predictors of memory encoding and consolidation. Neurosci Biobehav Rev 50:128–142.

Coltheart M (1981) The MRC psycholinguistic database. Q J Exp Psychol A 33:497–505.

Davachi L, Wagner AD (2002) Hippocampal contributions to episodic encoding: insights from relational and item-based learning. J Neurophysiol 88:982–990.

DuBrow S, Davachi L (2016) Temporal binding within and across events. Neurobiol Learn Mem 134:107–114.

Galli G (2014) What makes deeply encoded items memorable? Insights into the levels of processing framework from neuroimaging and neuromodulation. Front Psychiatr 5:1–8.

Galli G, Choy TL, Otten LJ (2012) Prestimulus brain activity predicts primacy in list learning. Cogn Neurosci 3:160–167.

Galli G, Feurra M, Pavone EF, Sirota M, Rossi S (2017) Dynamic changes in prefrontal cortex involvement during verbal episodic memory formation. Biol Psychol 12:536–544.

Galli G, Gebert D, Otten LJ (2013) Attentional resources modulate prestimulus brain activity related to memory encoding. Cortex 49:2239–2248.

Galli G, Griffiths V, Otten LJ (2014) Emotion regulation modulates anticipatory brain activity that predicts emotional memory encoding in women. Soc Cogn Affect Neurosci 9:378–384.

Galli G, Wolpe N, Otten LJ (2011) Sex differences in the use of anticipatory brain activity to encode emotional events. J Neurosci 31:12364–12370.

Heusser AC, Ezzyat Y, Shiff I, Davachi L (2018). Perceptual boundaries cause mnemonic trade-offs between local boundary processing and across-trial associative binding. J Exp Psychol Learn Mem Cogn 44:1075–1090.

Horner AJ, Bisby JA, Wang A, Bogus K, Burgess N (2016). The role of spatial boundaries in shaping long-term event representations. Cognition 154:151–164.

Kim H (2011) Neural activity that predicts subsequent memory and forgetting: a meta-analysis of 74 fMRI studies. Neuroimage 54:2446–2461.

Kučera H, Francis WN (1967) Computational analysis of present-day American English. Providence, RI: pnBrown University Press.

Li X, Nahas Z, Kozel FA, Anderson B, Bohning DE, George MS (2004) Acute left prefrontal transcranial magnetic stimulation in depressed patients is associated with immediately increased activity in prefrontal cortical as well as subcortical regions. Biol Psychol 55:882–890.

Machizawa MG, Kalla R, Walsh V, Otten LJ (2010) The time course of ventrolateral prefrontal cortex involvement in memory formation. J Neurophysiol 103:1569–1579.

Mangels JA, Picton TW, Craik FI (2001) Attention and successful episodic encoding: an event-related potential study. Cogn Brain Res 11:77–95.

van de Nieuwenhuijzen ME, van den Borne EW, Jensen O, van Gerven MA (2016). Spatiotemporal Dynamics of Cortical Representations during and after Stimulus Presentation. Front Syst Neurosci 9:10:42.

Otten LJ, Quayle AH, Akram S, Ditewig TA, Rugg MD (2006) Brain activity before an event predicts later recollection. Nat Neurosci 9:489–491.

Park H, Rugg, MD (2010) Prestimulus hippocampal activity predicts later recollection. Hippocampus 20:24–28.

R Core Team (2019). R: A language and environment for statistical computing. R Foundation for Statistical Computing. Vienna, Austria. URL https://www.R-project.org/.

Rossi S, Hallett M, Rossini PM, Pascual-Leone A (2011) Screening questionnaire before TMS: an update. Clin Neurophysiol 122:1686.

Rossi S, Innocenti I, Polizzotto NR, Feurra M, De Capua A, Ulivelli, M (2011) Temporal dynamics of memory trace formation in the human prefrontal cortex. Cereb Cortex 21:368–373.

Rossini PM, Burke D, Chen R, Cohen LG, Daskalakis Z, Di Iorio R, Di Lazzaro V, Ferreri F, Fitzgerald PB, George MS, Hallett M (2015) Non-invasive electrical and magnetic stimulation of the brain, spinal cord, roots and peripheral nerves: basic principles and procedures for routine clinical and research application. An updated report from an IFCN Committee. Clin Neurophysiol 126:1071–1107.

Silva M, Baldassano C, Fuentemilla L (2019) Rapid memory reactivation at movie event boundaries promotes episodic encoding. J Neurosci 0369–19.

Snodgrass JG, Corwin J (1988) Pragmatics of measuring recognition memory: applications to dementia and amnesia. J Exp Psychol Gen 117:34–50.

Sols I, DuBrow S, Davachi L, Fuentemilla L (2017) Event boundaries trigger rapid memory reinstatement of the prior events to promote their representation in long-term memory. Curr Biol 27:3499–504.

Staresina BP, Davachi L (2006) Differential encoding mechanisms for subsequent associative recognition and free recall. J Neurosci 26:9162–9172.

Viechtbauer, W. (2010). Conducting Meta-Analyses in R with the metafor Package. J Stat Softw 36:1–48.

